# A correlational study of ABCA3 and SCN4B as exercise-related biomarkers of patients with Stanford type A aortic dissection

**DOI:** 10.64898/2026.04.09.717394

**Authors:** Shuai Qiao, Tienan Chen, Bingxin Xie, Yongyong Han, Bin Wang, Yuanyuan Li, Beibei Jia, Naishi Wu

## Abstract

**Background:** Accumulating evidence indicates that moderate exercise may reduce the incidence of Stanford type A aortic dissection (TAAD), but the specific mechanisms remain unclear. This study aims to identify exercise-related biomarkers in TAAD patients and to investigate their underlying mechanisms.

**Methods:** Transcriptome data related to TAAD and exercise-related genes were obtained from publicly available databases. Candidate biomarkers for TAAD were identified through an integrative approach incorporating differential expression analysis, machine learning, and expression level assessment, leading to the construction of a diagnostic model. Subsequently, functional enrichment, immune infiltration, regulatory network analysis, and computational drug prediction were conducted to systematically investigate the pathological mechanisms and translational potential of the indentified biomarkers.

**Results:** ABCA3 and SCN4B were identified as exercise-related biomarkers in TAAD progression. A nomogram incorporating these two biomarkers exhibited strong diagnostic performance for identifying the disease. Functional enrichment analysis revealed potential involvement of these biomarkers in disease progression through pathways including circadian rhythm regulation and ribosome biosynthesis. Additionally, immune cells like M1 macrophages and naive B cells, as well as regulatory factors including hsa-miR-1343-3p and XIST, were found to be involved in this process. Finally, zonisamide and MRS1097 were identified through computation prediction as potential therapeutic drugs.

**Conclusion:** ABCA3 and SCN4B were identified as exercise-related biomarkers associatied with TAAD and represent potential valuable targets for both diagnosis and treatment strategies.

## 1. Introduction

Aortic dissection (AD) is a life-threatening cardiovascular disease associated with extremely high mortality rate, often described as “ticking time bomb” in human body. This condition occurs when the aorta intima develops a tear, allowing high-speed blood flow to dissect between the intima and media layers, resulting in the separation of the middle layer of the arterial wall along the long axis and forming true and false lumens. According to the Stanford classification, AD is categorized into type A and B based on the anatomical location of the entry tear and the dissection extent. Stanford type A aortic dissection (TAAD), the more lethal subtype, invariably involves the ascending aorta irrespective of the tear location [1, 2].The incidence of TAAD is increasing globally, exhibiting sex-specific disparities, where males have a higher overall incidence compared to females [3]. Various factors such as geography, population demographics, and methodologies for data identification influence its incidence rates, with a high estimated global prevalence. Approximately 20%-30% of TAAD cases have genetic origins, while acquired factors such as hypertension and smoking play a significant role [4, 5]. In TAAD, mortality increased by 1% per hour after onset,with untreated patients facing > 50% mortality within 48 hours. Thus, early diagnosis, immediate intervention, and rigorous follow-up are critical for improving patients survival rate. Despite advances in diagnostic methods, surgical techniques, and perioperative management,TAAD outcomes remain poor, with in-hospital mortality rates persisting at 22% and operative mortality at 18%[6]. These data urgently suggest the imperative to explore the molecular pathogenesis underlying TAAD and to identify biomarkers capable of mitigating its progression.

Exercise is widely recognized as a crucial intervention and preventive measure for cardiovascular diseases. A precise scientific classification of exercise intensity is essential for accurately assessing its cardiovascular protective effects. According to the latest guidelines from the American Heart Association, exercise intensity can be distinctly categorized: Moderate-intensity exercise, such as brisk walking, is defined as activities that elevate the heart rate to 50-70% of the maximum heart rate or correspond to a metabolic equivalent of 3-6 METs. In contrast, high-intensity exercise encompasses activities that raise the heart rate to 70-85% of the maximum heart rate or exceed a metabolic equivalent of 6 METs[7].Exercises of varying intensities influence the cardiovascular system via distinct molecular mechanisms. Moderate-intensity aerobic exercise can provide a protective effect by modulating the expression of inflammatory factors and diminishing the extent of pyroptosis[8].High-intensity exercise training can improve the overall cardiovascular regulation by enhancing the dynamic Starling mechanism and the sensitivity of the dynamic and static cardiac reflexes[9].This clear classification of intensity and the differences in mechanisms provide a foundation for further investigation into their association with specific cardiovascular diseases. Additional research suggests that exercise may offer a protective effect against the onset and progression of aortic dissection, a severe cardiovascular condition. Current evidence indicates that exercise can mitigate the degradation of the extracellular matrix (ECM) and preserve the structural integrity of the aortic wall by modulating the expression of inflammation-related genes and genes involved in ECM metabolism, thereby reducing the risk of aortic dissection[10]. In the pathological context of lysyl oxidase inhibition, aerobic exercise has been shown to decrease mortality and enhance core pathological features, including extracellular matrix degradation, thoracic aortic aneurysm, and dissection[11–13]. However, the precise mechanism underlying exercise’s effects in aortic dissection remain incompletely understood, and there are no studies to explore the biological function of exercise-related genes (ERGs) in TAAD. Elucidating these functions could provide new insights into disease mechanisms and identify promising new therapeutic targets.

This study employs a systematic transcriptomic analysis to establish a molecular link between exercise and TAAD. By integrating the differential expression profile of TAAD with a rigorously defined set of movement-related genes, machine learning algorithms are utilized to identify movement-related characteristic genes that exhibit differential expression in TAAD.Using a comprehensive approach that included experimental validation, functional enrichment analysis, immune microenvironment assessment, molecular regulatory network establishment, and drug prediction, the intricate involvement of exercise-related biomarkers in the pathogenesis of TAAD was systematically elucidated. These findings not only offer novel insights into therapeutic target identification for TAAD and establish a theoretical framework for understanding how moderate-intensity exercise affects TAAD at the molecular level. This research has important implications for developing personalized strategies to prevent and manage TAAD.

## 2. Materials and methods

### 2.1 Data source

A total of 2 transcriptome datasets (GSE153434 and GSE52093) linked to Stanford type A aortic dissection (TAAD) were obtained from Gene Expression Omnibus (GEO) database (https://www.ncbi.nlm.nih.gov/geo/). The details of these datasets were provided in **Table 1**. In this study, disease samples in each dataset were defined as case (TAAD) category, whereas the normal samples were designated as control category. In order to identify genes that are associated with exercise and may play a role in TAAD, we searched gene sets from MSigDB (https://www.gsea-msigdb.org/gsea/msigdb), specifically, using “exercise” as the keyword, and selecting species of Homo sapiens, and three related gene sets were identified: HP_ABNORMAL_CARDIAC_EXERCISE_STRESS_TEST, HP_ELEVATED_CREATINE_KINASE_AFTER_EXERCISE, and HP_EXERCISE_INTOLERANCE. The genes in these three gene sets were combined and duplicates were removed, resulting in 176 exercise-related genes listed in **S1 Table**. It should be noted that “exercise-related genes” is a broad functional classification based on phenotypic associations. The purpose of introducing this gene set is to prioritize differentially expressed genes (DEGs) that are biologically linked to exercise, a known protective intervention against cardiovascular diseases (CVD), in subsequent analyses, thus providing clues to potential biomarkers suggestive of physiological interventions.

**Table 1.** details of datasets.

| GSE series | Sample type | Sample size |  | Platform | RNA-sequencing<br>(RNA-seq) type |
| --- | --- | --- | --- | --- | --- |
|  |  | Case | Control |  |  |
| GSE153434 | Ascending aorta | 10 | 10 | GPL20795 | Bulk |
| GSE52093 | smooth muscle | 7 | 5 | GPL20795 |  |

### 2.2 Acquisition of exercise-associated differentially expressed genes

GSE153434 dataset was processed via DESeq (v 4.10.2) [14]to detect DEGs between TAAD and control categories. Based on the negative binomial distribution model, this tool can effectively correct for library size differences and gene expression heterogeneity in RNA-seq data, and is suitable for difference analysis of bulk transcriptome data. The Benjamini-Hochberg (BH) method was used to correct for multiple testing, and the screening criteria were |log2 Fold Change (FC)| > 0.5 and the adjust.P < 0.05. Next, the top 5 up- and down-regulated DEGs sorted by absolute log_2_ FC were labeled in volcano map, which was performed via ggplot2 (v 3.5.1) [15], and the top 40 DEGs were visualized via heatmap.plus (v 1.3) [16]. The genes that intersected between DEGs and ERGs were subsequently identified as DE-ERGs and plotted as venn diagrams with ggvenn (v 0.1.10) [17].

### 2.3 Functional enrichment

With ClusterProfiler (v 4.7.1.3) [18] and org.Hs.eg.db [19], Kyoto Encyclopedia of Genes and Genomes (KEGG) and Gene Ontology (GO) analyses were carried out to investigate biological processes and roles involved in DE-ERGs (*P* < 0.05). ClusterProfiler supports the integration of data from multiple sources (e.g., UCSC, Ensembl, NCBI, STRING, GENCODE, etc.), which facilitates the acquisition of functional annotations [20]. In the meantime, we carried out PPI network design to comprehend the relationships among DE-ERGs. In short, PPI network was built using DE-ERGs as input data and an interaction score > 0.4 (i.e., deleting isolated targets) according to Search Tool for the Retrieval of Interacting Genes (STRING) database (http://www/string-db.org/). The proteins with high-quality interactions were visualized by Cytoscape software (v 3.10.1) [21].

### 2.4 Machine learning and validation of biomarkers

To locate exercise-associated biomarkers in TAAD, DE-ERGs were evaluated using machine learning. First, Least Absolute Shrinkage and Selection Operator (LASSO) regression was performed for analysis. As a regularized linear model, LASSO incorporates an L1 penalty term (the sum of the absolute values of regression coefficients) to shrink the coefficients of irrelevant variables to zero, thereby achieving feature selection and preventing overfitting. In specific operations, we utilized the glmnet (v 4.1.8) [22]to construct a model under the binary classification setting (family = “binomial”), with the response variable being sample grouping (TAAD vs. control). The optimal penalty parameter λ (lambda.min) was determined via 5-fold cross-validation (type.measure = “deviance”), and genes with non-zero coefficients at this λ value were selected as LASSO feature genes. Second, the support vector machine-recursive feature elimination (SVM-RFE) algorithm was applied. Based on the support vector machine (SVM) model, this algorithm evaluates feature importance and recursively eliminates the least important features to identify the optimal feature subset. The analysis was performed using the caret (v 6.0.94) [23].To reduce redundancy, highly correlated features (correlation coefficient > 0.9) were removed in advance. An SVM model with a radial basis function kernel (method = “svmRadial”) was adopted, and 5-fold cross-validation (method = “cv”) was used to evaluate the model performance of feature subsets with different sizes. Finally, the feature set corresponding to the lowest error rate was selected as the SVM-RFE feature genes. Third, the Boruta algorithm was employed for comprehensive feature importance assessment. Based on random forest, this algorithm generates randomly permuted “shadow features” for each real feature and compares the importance of real features with that of the optimal shadow features (measured by Z-scores), thereby determining whether a feature is statistically significant. The analysis was conducted using the Boruta (v 8.0.0) [24], with the significance level set at pValue = 0.001. Iterations were performed until all features were clearly classified as “Confirmed”, “Tentative”, or “Rejected”, and genes confirmed as important were included in the Boruta feature genes. Finally, the intersection of three feature genes was used to determine the candidate genes. Furthermore, the expression levels of candidate genes within GSE153434 and GSE52093 were subjected to Wilcoxon test (P < 0.05), and the outcomes were illustrated with violin plots. Genes with consistent expression trends and significant expression differences between groups across all datasets were ultimately defined as biomarkers.

### 2.5 Nomogram establishment and validation

To assess the diagnostic value of biomarkers related to exercise in TAAD, nomogram model integrating biomarkers was developed by rms package (v 6.8.1) [25] within GSE153434 dataset. Using this nomogram model, the probabilities of being diagnosed with TAAD could be predicted. Moreover, pROC (v 1.18.5) [26], rms (v 6.8.1) [25], and ggDCA packages (v 1.1) [27]. Assessment metrics included , which required an area under the curve (AUC) exceeding 0.7, a calibration curve assessed via the Hosmer-Lemeshow (HL) test with a significance level set at p > 0.05, and decision curve analysis (DCA) to gauge the clinical utility.

### 2.6 Gene set enrichment analysis (GSEA)

The association between biomarkers and the other genes was evaluated independently in GSE153434 dataset and sorted by correlation coefficient from biggest to lowest. Next, based on the reference gene set “c2.cp.v7.2.symbols.gmt” from MSigDB, Gene Set Enrichment Analysis (GSEA) was performed on the ranked gene list using the clusterProfiler (v 4.7.1.3)package. This analytical method evaluates the degree of enrichment of a gene set at the top or bottom of a phenotype-related gene list by calculating the weighted enrichment score (WES) of the gene set. In the analysis, the Benjamini-Hochberg (BH) method was used to correct for multiple testing, with the filtering criteria set as (|normalized enrichment score (NES)| > 1, adjust.*P* < 0.05). Subsequently, in all TAAD samples of training set, the associations between biomarkers and genes in the shared pathways were further explored. Specifically, “cor” function (p<0.05) was first used to analyze the correlations between each biomarker and genes in the circadian rhythm pathway via Spearman test. Thereafter, the same function was employed to assess the correlations between circadian rhythm pathway scores and each biomarker.

### 2.7 Immune microenvironment analysis

Within the GSE153434 dataset, we utilized the CIBERSORT algorithm to investigate the differences in the immune microenvironment between the TAAD group and the control group. Based on deconvolution calculations of the gene expression signature matrix, this algorithm estimates the relative infiltration proportions of 22 immune cell types in mixed tissue samples. First, we applied this algorithm to separately visualize the infiltration abundances of various immune cell types in the samples from the two groups. Subsequently, differential immune infiltrating cell types were recognized by Wilcoxon test (P < 0.05). Thereafter, psych (v 2.4.3) [28] was employed to perform Spearman correlation analysis, which correlated differential immune infiltrating cell types with each other and with biomarkers (|cor| > 0.3, p < 0.05).

### 2.8 Single-cell expression profile analysis of target genes

To elucidate the potential association between biomarkers and immune cells, single-cell RNA expression data for the two genes were obtained from the Human Protein Atlas (HPA) database (https://www.proteinatlas.org/). The expression information of the target genes in different human single-cell types was retrieved using the HPA open API, and the normalized expression level nCPM (normalized protein-coding transcripts per million) was extracted as the expression indicator. The nCPM values of the identified biomarkers in each single-cell type were sorted in descending order, and the top 10 cell types with the highest expression levels were selected for subsequent analysis. Visualization of the top 10 highly expressed cell types and their corresponding expression levels was performed using the ggplot2 (v 3.5.1) and patchwork (v 1.2.0) packages in R software to evaluate the specific expression distribution characteristics of the target genes across different single-cell types, thereby providing evidence for determining whether the genes were expressed by immune cells.

### 2.9 Molecular regulatory networks

To investigate the molecular regulatory mechanisms of biomarkers, microRNAs (miRNAs) targeting them were initially predicted using microRNA target prediction database (miRDB, http://mirdb.org) and encyclopedia of RNA interactomes (ENCORI/starBase) database (http://starbase.sysu.edu.cn/), respectively. The shared miRNAs predicted from both databases were utilized as key miRNAs and were used to separately predict long non-coding RNAs (lncRNA)s in ENCORI database. Subsequently, lncRNA-miRNA-biomaker network was built using Cytoscape (v 3.10.1) [29], based on lncRNAs, key miRNAs, and biomarkers.

### 2.10 Drug prediction and molecular docking

In order to explore potential drugs for treating TAAD, Drug-Gene Interaction Database (DGIdb) (http://www.dgidb.org/) was employed to search for potential drugs. The relationships between the biomarkers and potential drugs were visualized via Cytoscape software (v 3.10.1). Then, the 3D molecular structures of drugs (with the highest interaction score) and the 3D structure of biomarkers were downloaded from PubChem database (https://pubchem.ncbi.nlm.nih.gov/) and Protein Data Bank (PDB, https://www.rcsb.org/), respectively. The binding sites of the target protein were analyzed to identify the corresponding docking active pockets, which were then validated through 50 independent docking simulations using AutoDock (v 4.2) [30]. Theoretically, a binding energy lower than -5 kcal/mol between the receptor protein and ligand compound indicates a strong binding affinity.

### 2.11 Reverse transcription polymerase chain reaction (RT-qPCR)

To preliminarily validate the key genes screened by bioinformatics analysis, this study collected aortic dissection tissue samples from five TAAD patients and five healthy individuals at the Second Hospital of Tianjin Medical University. This research was approved by the Ethics Committee of the Second Hospital of Tianjin Medical University (Approval No: KY2025K248), and all patients signed informed consent forms. First, 50 mg of tissue from each sample was mixed with 1 ml of TRIzol (Ambion, USA) to ensure full homogenization and grinding. After standing on ice for 10 minutes, 0.2 ml of chloroform was added to extract the RNA aqueous phase. An equivalent volume of cold isopropanol was then added to extract the RNA. Following further RNA measurement, reverse transcription was initiated immediately. The reaction system for DNA synthesis was set up according to the manufacturer’s instructions for the Hifair® Ⅲ 1st Strand cDNA Synthesis SuperMix for qPCR (Yeasen, 11141ES60, Shanghai, China) and SwsScript All-in-One First-strand-cDNA-synthesis SuperMIx for qpcr (One step gDNA Remover) was conducted on ordinary PCR machine (BIO-RAD, U.S.A) and XLFZ006 real-time fluorescence quantitative PCR equipment (BIO-RAD, U.S.A) at 40 cycles. **Table 2** presents primer sequences, and biomarker expression was assessed using the 2-ΔΔCt approach, employing GAPDH as a reference gene for normalization [31]. Finally, Graphpad Prism (v 10.1.2) [32] was employed to plot and calculate the p value. This experiment aimed to preliminarily validate the expression trends of the candidate biomarkers identified through bioinformatics analysis using independent clinical samples, thereby enhancing the reliability of the research findings.

**Table 2.**
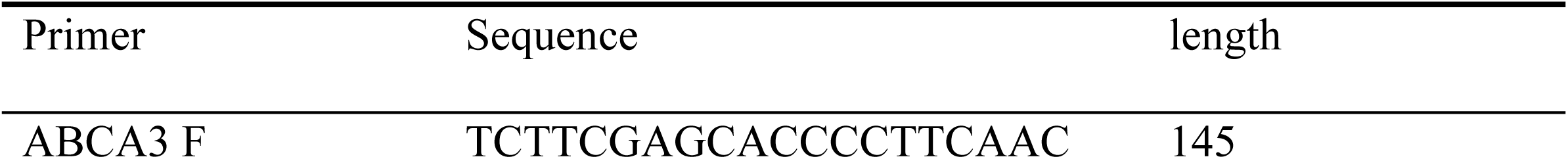
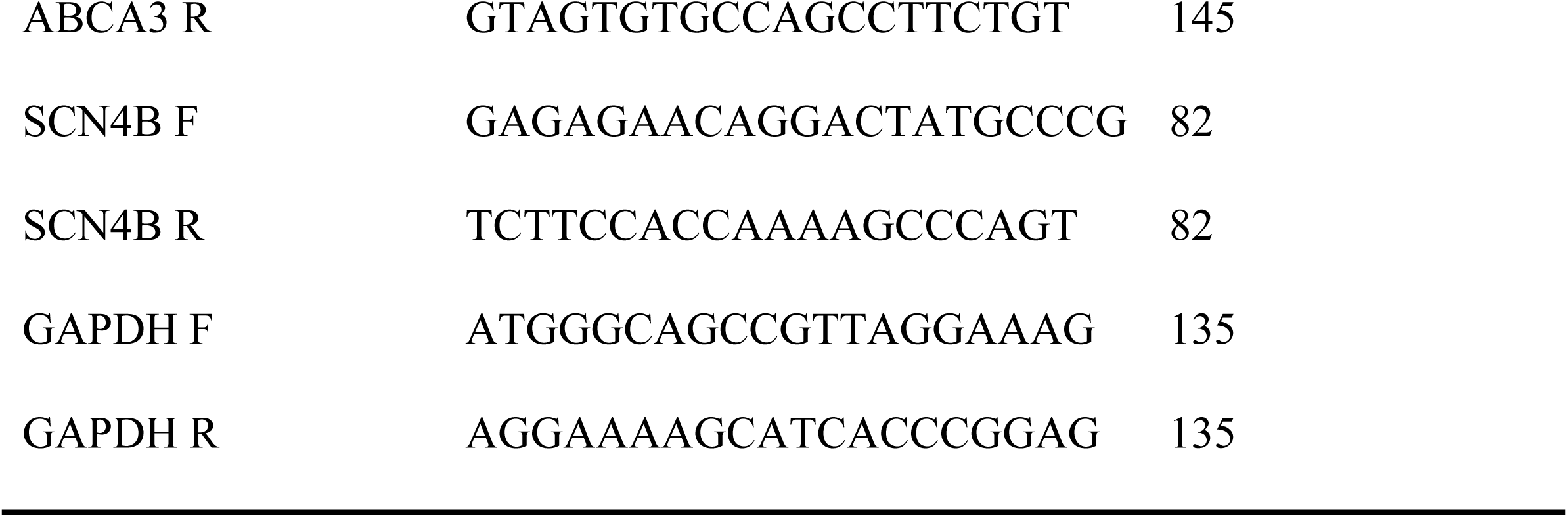
primer sequences.

### 2.12 Statistical analysis

Bioinformatics analyses were conducted using R (v 4.2.3). Differences between two groups were came true by Wilcoxon rank sum test, and differences between PCR experimental groups were obtained through t test. There was a statistically significance of *P* < 0.05.

## 3. Results

### 3.1 Various functional pathways and complex PPI network of DE-ERGs

Performing differential expression analysis between TAAD and controls in GSE153434, a total of 2,000 DEGs were determined, along with 1,259 decreased and 741 increased genes (**Fig 1a-b**). By crossing DEGs with ERGs, 17 DE-ERGs were obtained (**Fig 1c**). The future enrichment analyses of DE-ERGs revealed that ERGs in TAAD were primarily associated with structural maintenance and functional regulation of the cardiovascular system. Specifically, GO analysis showed that DE-ERGs were significantly enriched in 214 biological process (BP), 61 molecular function (MF), and 38 cellular component (CC) terms, involving actin-mediated cell contraction, T-tubule, and gated channel activity (**Fig 1d**). Additionally, these differentially expressed enhancer regions were highly enriched in 12 KEGG pathways, including myocardial contraction-related pathways (such as adrenergic signaling in cardiomyocytes, cardiac muscle contraction, hypertrophic cardiomyopathy pathways), energy metabolism pathways (glycolysis/gluconeogenesis, propanoate metabolism), and cytoskeletal regulation (cytoskeleton in muscle cells, motor proteins) (**Fig 1e**). PPI network revealed extensive interactions among 17 DE-EPRG proteins, with MYH6 exhibiting the strongest interactions with other genes. Notably, COL12A1 was found to interact exclusively with COL9A2 (**Fig 1f**). In summary, the enrichment analysis results highlight the complex interplay of various biological processes and molecular pathways in TAAD, emphasizing the importance of maintaining cardiovascular structure and function. These insights provide a foundation for further research into the pathogenesis of TAAD and the development of targeted diagnostic and therapeutic approaches.

**Fig 1.**
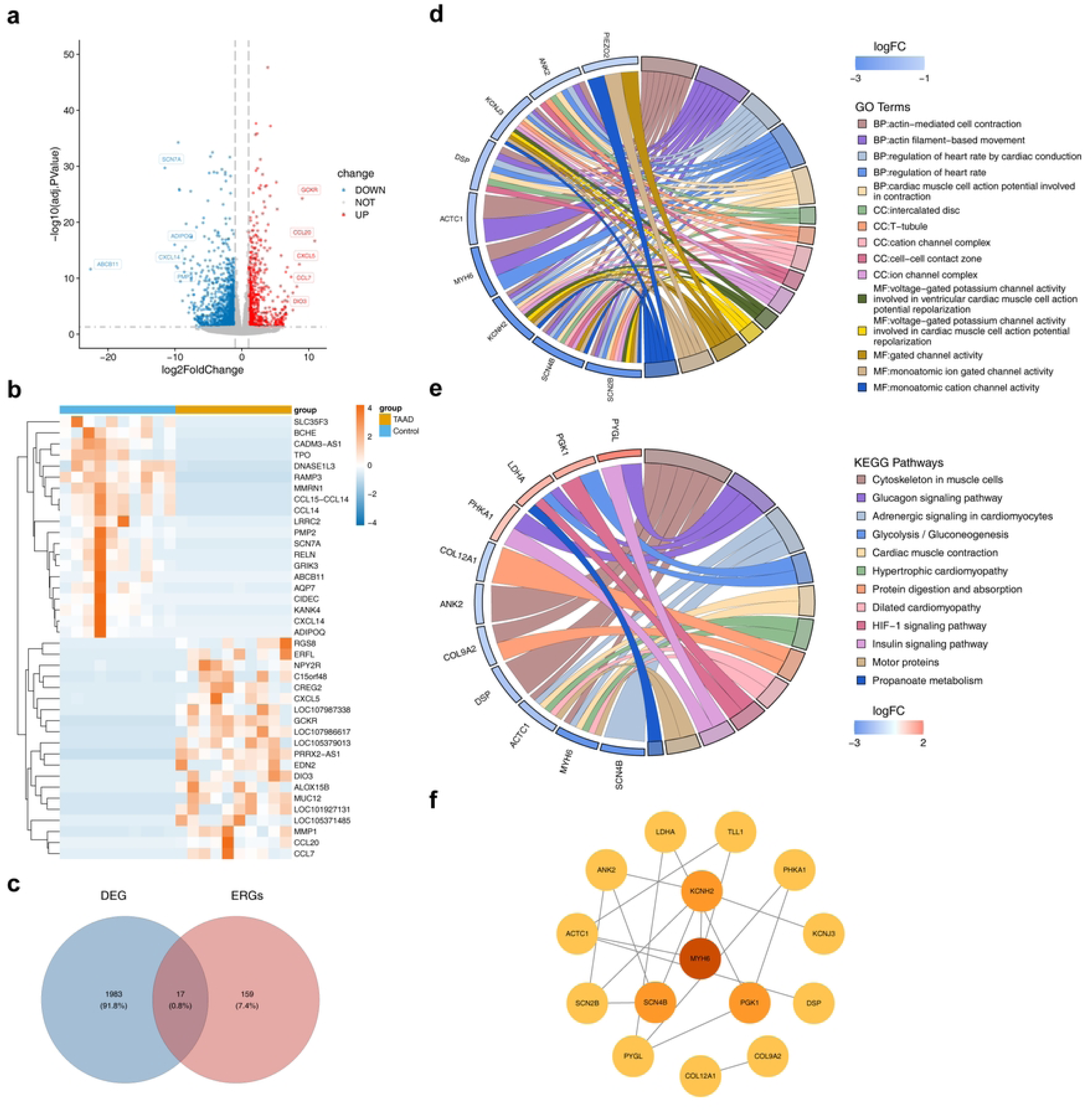
Various functional pathways and complex PPI network of DE-ERGs. (a-b) Performing differential expression analysis between TAAD and controls. (c) By crossing DEGs with ERGs, 17 DE-ERGs were obtained. (d) GO analysis showed that DE-ERGs were significantly enriched in 214 biological process (BP), 61 molecular function (MF), and 38 cellular component (CC) terms. (e) Differentially expressed enhancer regions were highly enriched in 12 KEGG pathways. (f) PPI network revealed extensive interactions among 17 DE-EPRG proteins, with MYH6 exhibiting the strongest interactions with other genes.

### 3.2 ABCA3 and SCN4B were recognized as biomarkers of TAAD

When lambada.min = 0.004366, 5 LASSO feature genes were selected : ABCA3, LDHA, PGK1, PHKA1, SCN4B (**Fig 2a**). With respect to 17 DE-ERGs evaluated with SVM-RFE, the results suggested that when the SVM-RFE model had the lowest error rate, it corresponded to the 5 feature genes were SCN4B, PGK1, PHKA1, SCN2B, and ABCA3 (**Fig 2b**). Meantime, a total of 13 feature genes were selected via the Boruta algorithm (**Fig 2c**). After overlapping three sets of feature genes, 4 candidate genes were obtained (ABCA3, PGK1, PHKA1, SCN4B) (**Fig 2d**). The expression profiles of signature genes were further analyzed. It was noted that ABCA3 and SCN4B exhibited consistent expression trends and significant expression differences between case and control groups (*P* < 0.05) across all TAAD-related (GSE153434 and GSE52093) datasets (**Fig 2e**). Consequently, ABCA3 and SCN4B were identified as biomarkers related to exercise in TAAD. Notably, in clinical samples, the expression levels of both ABCA3 and SCN4B were decreased in TAAD (**Fig 2f**), which was consistent with the bioinformatics results. This suggests that the loss of their functions may be associated with disease progression and warrants further investigation.

**Fig 2.**
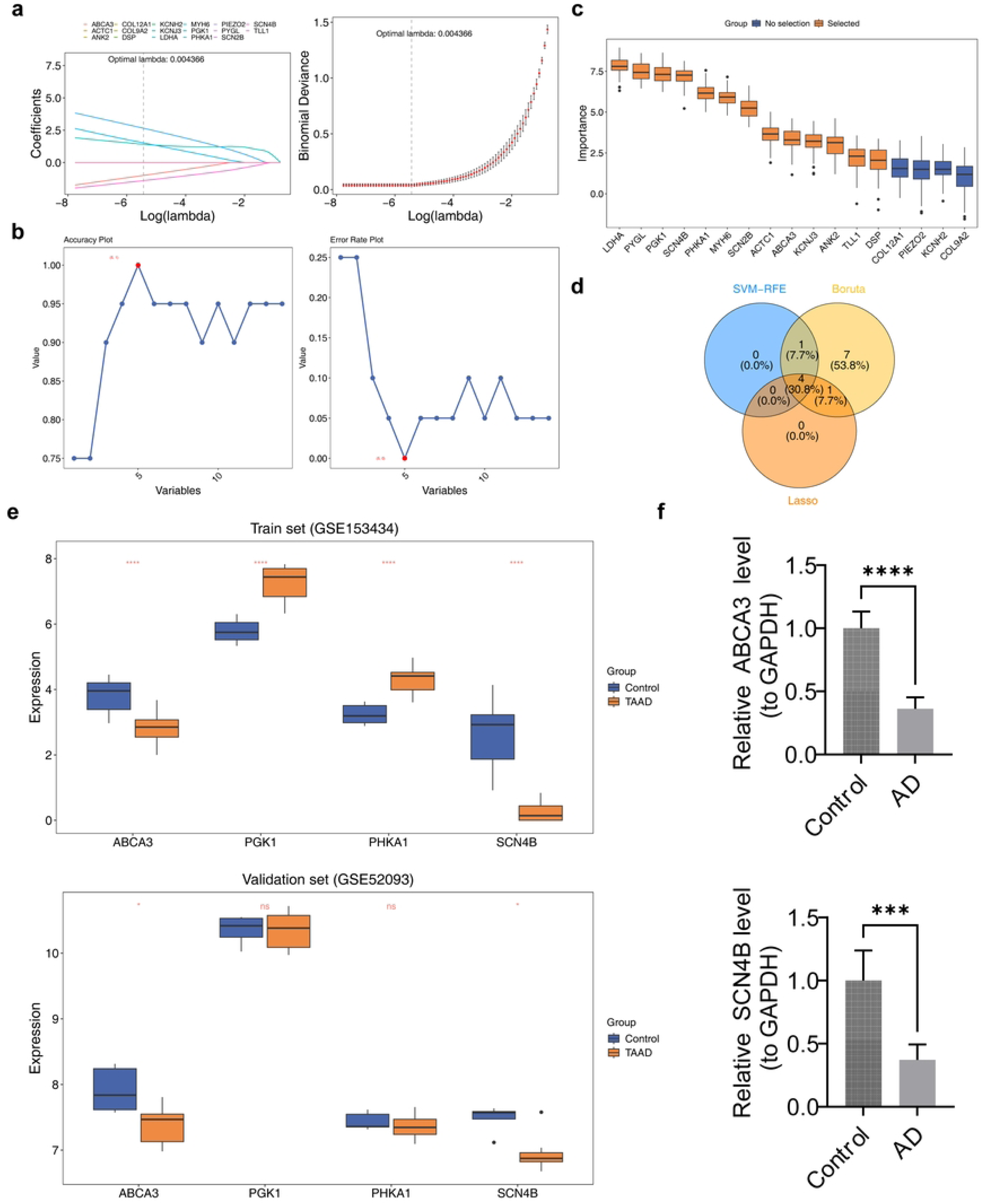
ABCA3 and SCN4B were recognized as biomarkers of TAAD. (a) When lambada.min = 0.004366, 5 LASSO feature genes were selected : ABCA3, LDHA, PGK1, PHKA1, SCN4B. (b) When the SVM-RFE model had the lowest error rate, it corresponded to the 5 feature genes were SCN4B, PGK1, PHKA1, SCN2B, and ABCA3. (c) A total of 13 feature genes were selected via the Boruta algorithm. (d) Overlapping three sets of feature genes, 4 candidate genes were obtained (ABCA3, PGK1, PHKA1, SCN4B). (e) ABCA3 and SCN4B exhibited consistent expression trends and significant expression differences between case and control groups (P < 0.05) across all TAAD-related datasets. (f) In clinical samples, the expression levels of both ABCA3 and SCN4B were decreased in TAAD (P < 0.05).

### 3.3 Outstanding diagnostic value and crucial functional pathways of biomarkers

Based on biomarkers, nomogram models for TAAD were established within GSE153434 dataset (**Fig 3a**). It was observed that higher total points were associated with greater probabilities of being diagnosed with TAAD. Importantly, the AUC value in ROC curves was 0.990 (**Fig 3b**), the slopes of the corresponding calibration curves approached 1 (HL test, *P* = 0.996) (**Fig 3c**), and the overall net benefit of the nomogram was higher than that of individual genes (**Fig 3d**). These results indicate that the nomogram model demonstrates potential as an effective predictive tool in clinical decision-making for TAAD, with ABCA3 and SCN4B being of particular diagnostic value for the disease.

**Fig 3.**
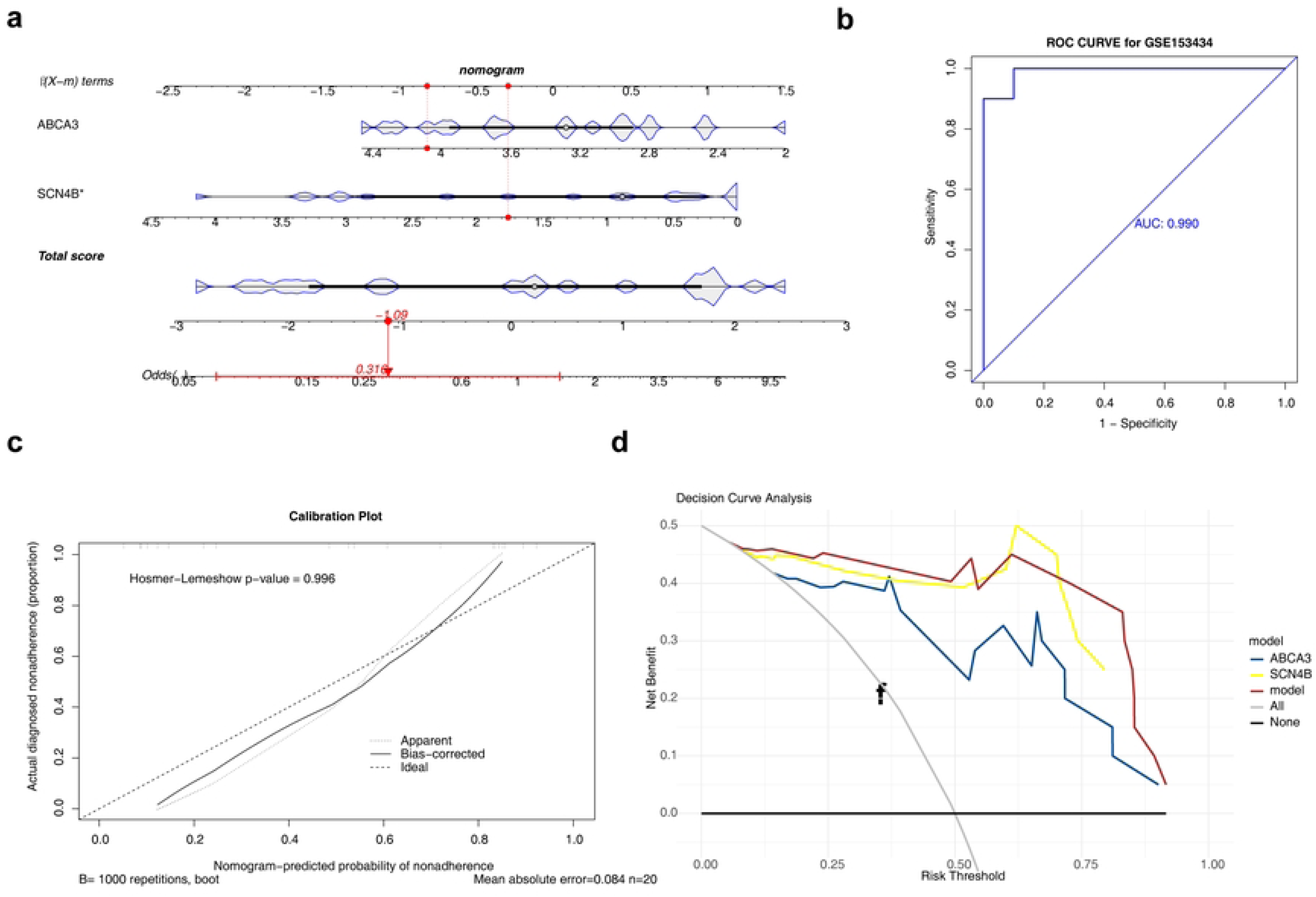
Diagnostic value of biomarkers. (a) Nomogram models for TAAD were established within GSE153434 dataset. (b) The AUC value in ROC curves was 0.990.(c) The slopes of the corresponding calibration curves approached 1 (HL test, *P* = 0.996). (d)The overall net benefit of the nomogram was higher than that of individual genes.

In the GSE153434 dataset, the functional pathways of biomarkers were explored. ABCA3 and SCN4B showed relatively similar functions, both being associated with “circadian entrainment”, “ribosome biogenesis in eukaryotes”, and “ribosome” (adjusted P < 0.05) (**Fig 4a-b**). Circadian rhythms influence multiple physiological processes in the body, such as blood pressure, heart rate, and hormone levels, all of which were closely related to cardiovascular health. ABCA3 was significantly positively correlated with JMJD7 - PLA2G4B in the circadian rhythm pathway (cor > 0.85, *P* < 0.05), while SCN4B was significantly negatively correlated with GNB4 (cor < 0.72, *P* < 0.05) (**Fig 4c-d**). Overall, both ABCA3 and SCN4B were positively correlated with the GSEA Scores of the circadian rhythm pathway (**Fig 4e-f**), indicating that as the pathway activity (GSEA scores increase), the expression levels of these two genes tend to increase.

**Fig 4.**
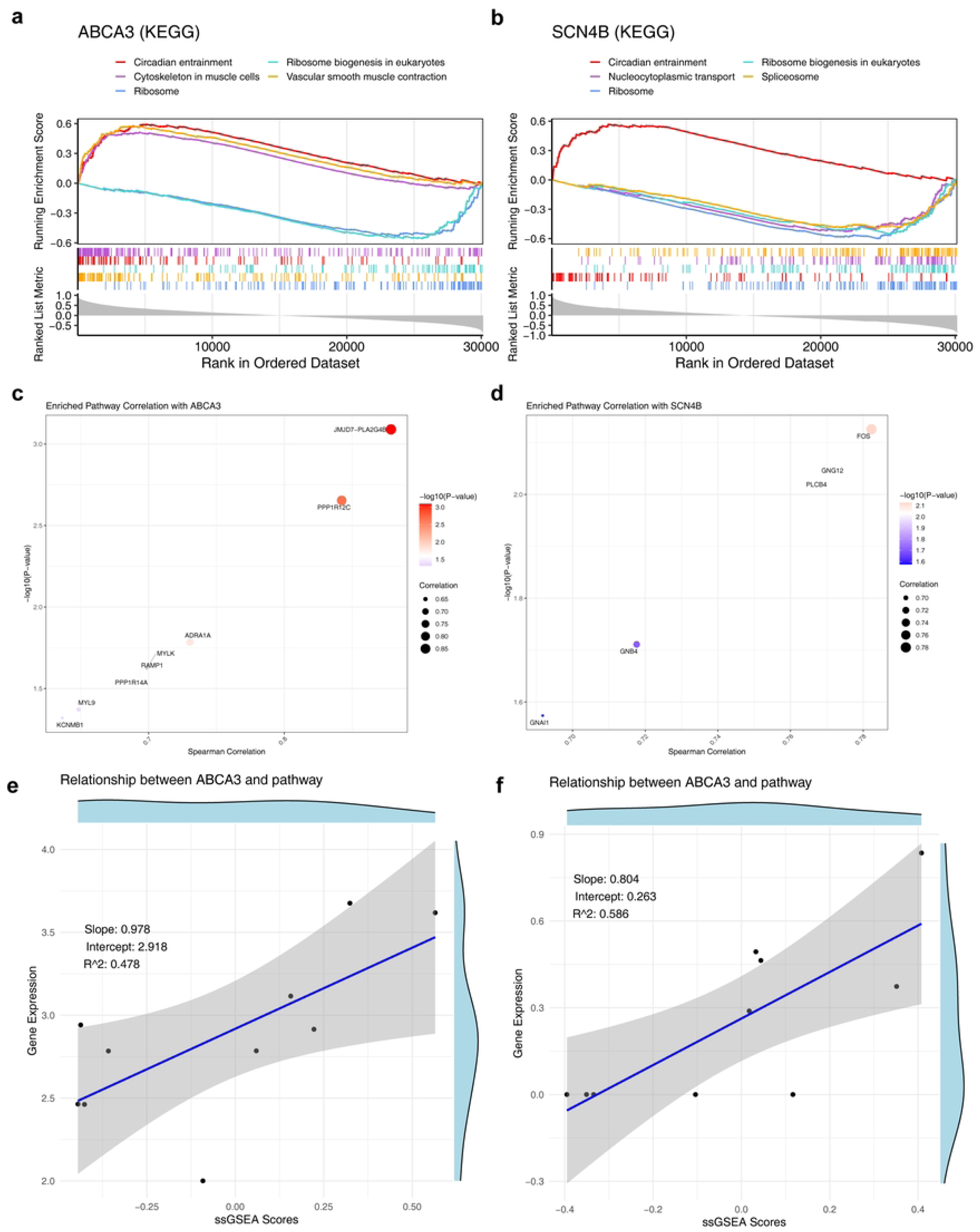
The functional pathways of biomarkers in the GSE153434 dataset. (a-b) ABCA3 and SCN4B showed relatively similar functions. (c-d) ABCA3 was significantly positively correlated with JMJD7 - PLA2G4B in the circadian rhythm pathway (cor > 0.85, *P* < 0.05), while SCN4B was significantly negatively correlated with GNB4 (cor < 0.72, *P* < 0.05). (e-f) Both ABCA3 and SCN4B were positively correlated with the GSEA Scores of the circadian rhythm pathway.

### 3.4 Immune-related activities could have a crucial role in TAAD

The infiltration ratios of 22 types of immune - infiltrating cells between the TAAD and control categories in GSE153434 dataset were described in detail. The infiltration level of M2 macrophages was relatively high in the samples (**Fig 5a**). Notably, the infiltration levels of activated mast cells, M0 macrophages, neutrophils, and monocytes were significantly increased in the TAAD category (*P* < 0.05), while those of M1 macrophages, naive B cells, and gamma delta T cells were significantly decreased (*P* < 0.05) (**Fig 5b**). Moreover, the study found that both neutrophils and monocytes were positively correlated with M0 macrophages (cor > 0.5). Similarly, naive B cells were negatively correlated with M0 macrophages, neutrophils, monocytes, and activated mast cells (cor < -0.5) (**Fig 5c**). ABCA3 was significantly negatively correlated with neutrophils (cor < 0.4, p < 0.001) and SCN4B was significantly positively correlated with naive B cells (cor > 0.6, p < 0.001) (**Fig 5d**). In conclusion, there were differences in the characteristics of the immune microenvironment between the TAAD and control groups, and these differences were significantly correlated with the locomotor genes.

**Fig 5.**
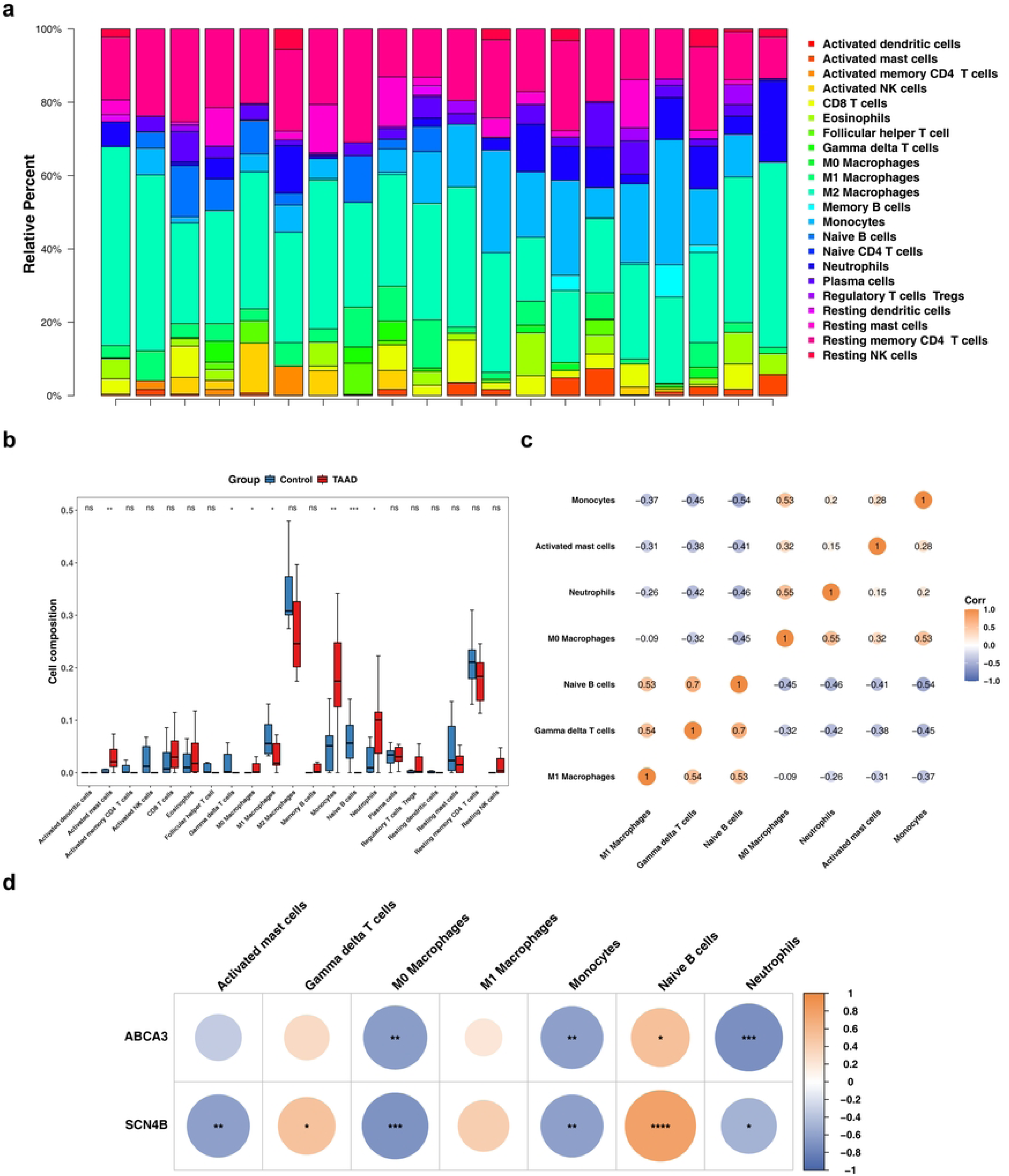
Immune-related activities have a crucial role in TAAD. (a) The infiltration ratios of 22 types of immune - infiltrating cells between the TAAD and control categories in GSE153434 dataset were described in detail. (b) The infiltration levels of activated mast cells, M0 macrophages, neutrophils, and monocytes were significantly increased in the TAAD category (*P* < 0.05), while those of M1 macrophages, naive B cells, and gamma delta T cells were significantly decreased (*P* < 0.05). (c) Both neutrophils and monocytes were positively correlated with M0 macrophages (cor > 0.5). Naive B cells were negatively correlated with M0 macrophages, neutrophils, monocytes, and activated mast cells (cor < -0.5). (d) ABCA3 was significantly negatively correlated with neutrophils (cor < 0.4, *P* < 0.001) and SCN4B was significantly positively correlated with naive B cells (cor > 0.6, *P* < 0.001)

### 3.5 Single-cell expression profile characteristics of ABCA3 and SCN4B

To explore the nature of the association between ABCA3 and SCN4B and immune cell infiltration, the single-cell expression profile characteristics of the two genes were analyzed using the HPA database. The results showed that ABCA3 and SCN4B exhibited significant expression specificity across different human single-cell types, but no immune cells were detected among the cell types with high expression of either gene (**Fig 6**). Specifically, the cell type with the highest expression of SCN4B was pericytes, with normalized expression level (nCPM) significantly higher than that of other cell types; the top 10 cell types with high expression also included vascular endothelial cells, smooth muscle cells, and other vascular wall-associated cells, while major immune cells such as macrophages, neutrophils, and T cells were absent. For ABCA3, the cell type with the highest expression was type II alveolar epithelial cells, and the top 10 cell types with high expression were all lung tissue-associated epithelial cells; similarly, no high expression signals were detected for immune cells such as macrophages or neutrophils. These findings indicated that ABCA3 and SCN4B were likely not highly expressed by the immune cells infiltrating the TAAD lesion tissues themselves, and their correlation with immune cell infiltration might rather reflect the interaction between the altered gene expression in non-immune cells of the aortic wall and the remodeling of the local immune microenvironment.

**Fig 6.**
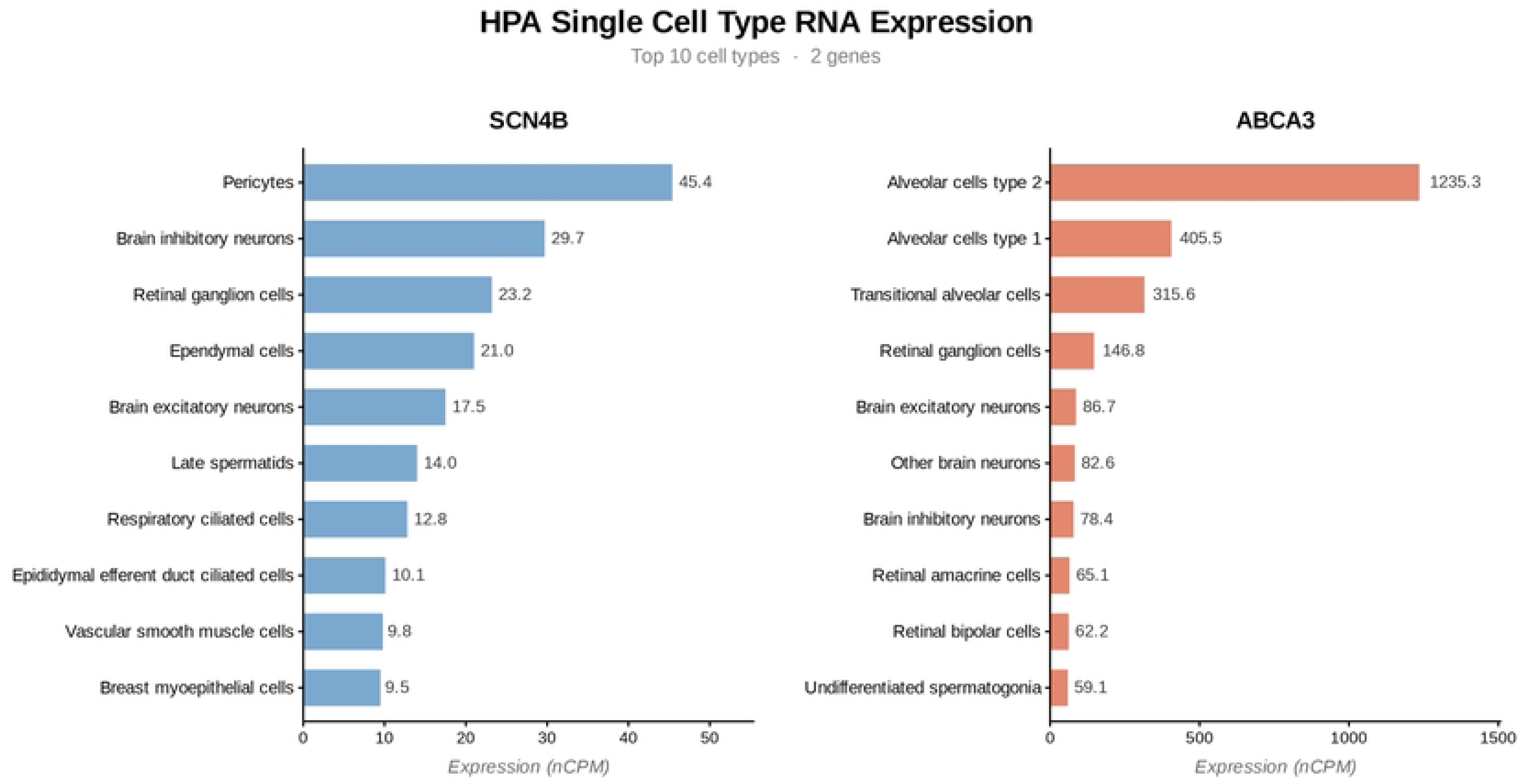
Expression of ABCA3 and SCN4B in different cell types.

### 3.6 A variety of molecules and drugs could target biomarkers in TAAD

Focusing on biomarkers, one key miRNA (hsa-miR-1343-3p) and 48 key lncRNAs (such as XIST) were found to be able to target and regulate ABCA3 according to miRDB. Additionally, three key miRNAs (e.g., hsa-miR-4770) and 63 key lncRNAs (e.g., MALAT1) were identified to target and regulate SCN4B (**Fig 7a**). The identified biomarkers, ABCA3 and SCN4B, are regulated by specific miRNAs and lncRNAs, indicating a complex post-transcriptional regulatory network that may play a role in the pathogenesis of TAAD.

**Fig 7.**
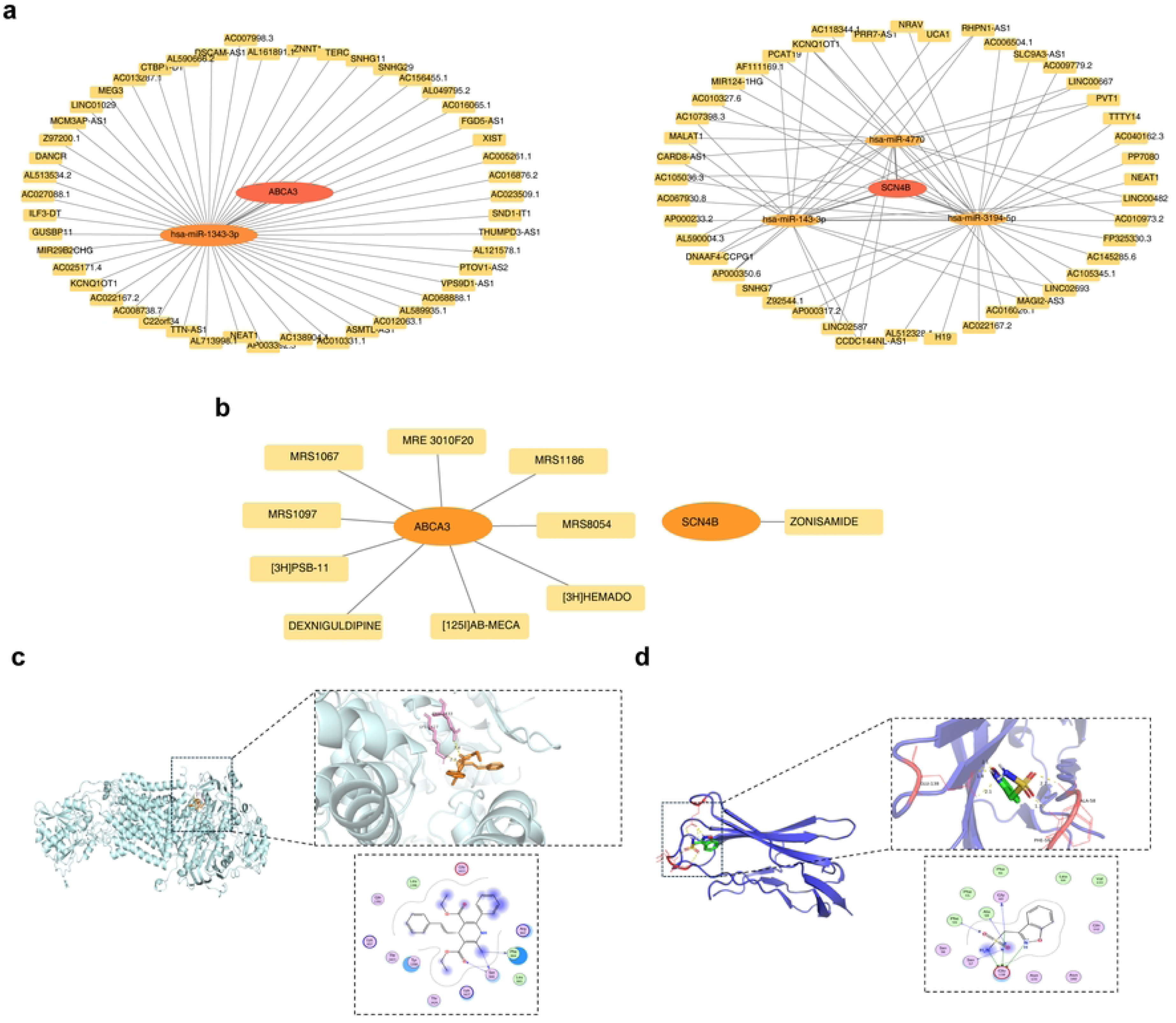
A variety of molecules and drugs could target biomarkers in TAAD. (a) The identified biomarkers, ABCA3 and SCN4B, are regulated by specific miRNAs and lncRNAs. (b) The DGIdb database predicted that nine drugs could bind to ABCA3, and one drug could bind to SCN4B. (c) Zonisamide binds to ABCA3 with a binding energy of -5.08 kcal/mol. This binding was stabilized by hydrogen bonds, hydrophobic interactions, and π - π stacking, involving residues such as Ser666 and Phe664. (d)Hydrogen bonds formed with Gly^60^ and Glu^138^ in the binding pocket contribute to the stable interaction between the small molecule and the protein.

The DGIdb database predicted that nine drugs could bind to ABCA3, and one drug could bind to SCN4B (**Fig 7b**). Zonisamide binds to ABCA3 with a binding energy of -5.08 kcal/mol. This binding was stabilized by hydrogen bonds, hydrophobic interactions, and π - π stacking, involving residues such as Ser666 and Phe664 (**Fig 7c**). The binding energy between SCN4B and MRS1097 was -7.3 kcal/mol. Hydrogen bonds formed with Gly^60^ and Glu^138^ in the binding pocket contribute to the stable interaction between the small molecule and the protein (**Fig 7d**). These results indicated that stable interactions may have been established between the studied proteins and the corresponding compounds, suggesting that these compounds could be potential ligands or drugs.

## 4. Discussion

TAAD is a devastating disease that requires a coordinated multidisciplinary approach for rapid diagnosis and treatment delivery. Surgical reconstruction for aortic aneurysm and dissection has been an important facet of the Stanford program[33]. Although immediate surgical intervention significantly increased survival rate after acute type A aortic dissection (ATAAD), operative mortality remained high[34]. Extensive medical literature has unequivocally established the significant and measurable advantages of exercise in the primary prevention of cardiovascular disease (CVD) [35, 36]. Furthermore, research has indicated that individuals with a high genetic predisposition to CVD who engage in regular physical activity exhibit a 31% lower incidence of TAAD compared to their sedentary counterparts [37], underscoring the protective effects of moderate aerobic exercise against TAAD. Nonetheless, the molecular underpinnings of how exercise influences the development of aortic dissection entail a complex regulatory network involving multiple pathways, with a dearth of systematic investigations elucidating the biological functions and regulatory mechanisms of ERGs in TAAD. Therefore, further investigation into the molecular pathogenesis of TAAD is warranted to identify biomarkers capable of mitigating disease progression. Through analysis of bulk RNA sequencing data, two exercise-related key genes (ABCA3 and SCN4B) were identified in TAAD using integrated bioinformatics approaches. Subsequent characterization of their diagnostic value and potential biological mechanisms has provided new insights into TAAD pathobiology, offering promising avenues for both diagnostic and therapeutic development.

Studies on cardiovascular structural and functional regulation have revealed that differentially expressed motion-related genes (DE-ERGs) are significantly enriched in actin-mediated cell contraction, T tube (transverse tube) structure formation and ion channel gating activity, which is highly consistent with the current understanding of the pathogenesis of TAAD. Recent studies [38] have demonstrated that vascular smooth muscle cell dysfunction as a key factor in aortic structural instability. This study suggests that DE-ERGs may impair aortic wall mechanical homeostasis by disrupting normal cardiac cycle dynamics through cytoskeletal regulation.

KEGG further revealed that DE-ERGs are significantly involved in key metabolic and signal pathways such as glycolysis/xenogenesis, myocardial contraction, the adrenergic signaling pathway and the hypertrophic cardiomyopathy (HCM) pathway. Under pathological conditions, abnormal energy metabolism may lead to insufficient ATP supply in cardiomyocytes, which in turn triggers abnormal activation of compensatory signaling pathways[39]. This vicious cycle accelerates vascular wall degeneration, while HCM pathway may induce cardiomyocyte pathological hypertrophy and interstitial fibrosis[40].

ABCA3 (ATP-binding cassette subfamily A member 3), an important member of the ATP-binding cassette transporter superfamily and is mainly expressed in alveolar type II epithelial cells [41]. This study found that the expression of this gene was significantly down-regulated in the aortic tissue of TAAD patients, which provides a new perspective for understanding its potential function in the cardiovascular system. Although direct research on ABCA3 in the cardiovascular system is relatively scarce at present, existing studies have shown that other members of the ABC transporter superfamily are widely present in vascular parietal cells and affect the cellular signal transduction process by regulating the lipid composition of the cell membrane, the integrity of lipid rafts, and the sorting and function of membrane proteins[42, 43].These transporters achieve substrate transport through a synergistic movement mechanism[44], and its abnormal function may lead to an imbalance in lipid metabolism within cells, thereby affecting the transduction of inflammatory signals in the local microenvironment [45]. Based on the findings of this study, a hypothesis can be proposed: The down-regulation of ABCA3 may be related to local lipid metabolism disorders and the activation of pro-inflammatory signals, and the inflammatory response has been regarded as an important driving factor for the occurrence and development of TAAD [46].In addition, abnormal function of the ABC transporter may interfere with the adhesion stability of vascular smooth muscle cells to the extracellular matrix, which is closely related to abnormal lipid rafts [47, 48]. Abnormalities in lipid rafts may weaken the adaptability of smooth muscle cells to mechanical loads and reduce the structural support of the extracellular matrix [49]. This suggests that the decreased expression of ABCA3 in aortic tissue may be involved in the potential mechanism of maintaining the integrity of the aortic membrane structure by affecting lipid homeostasis, cell adhesion and extracellular matrix balance. On the other hand, moderate exercise has been proven to have anti-inflammatory effects and can regulate lipid metabolism [50]. Therefore, we speculate that exercise may help maintain lipid homeostasis and inflammatory balance of the aortic wall by up-regulating the expression of ABCA3 or improving its function, thereby protecting the function of smooth muscle cells and delaying the degradation of extracellular matrix, which may play a protective role in the pathological process of TAAD. However, this mechanism hypothesis still needs to be further verified through vascular smooth muscle cell-specific ABCA3 knockout or overexpression models.

The SCN4B gene encodes the β4 subunit of the voltage-gated sodium channel (Navβ4), which is a key auxiliary subunit that regulates the function of the voltage-gated sodium channel (VGSC)[51]. Studies have shown that Navβ4 plays an important role in excitable tissues such as myocardium and neurons by regulating membrane expression, localization and electrophysiological properties of α subunit [52–54]. This finding has important clinical significance in that SCN4B expression is significantly down-regulated in aortic tissues of TAAD patients. Although research on sodium channels has mostly focused on excitable cells, recent studies have revealed that sodium channel complexes also exist in vascular smooth muscle cells and are involved in regulating intracellular calcium homeostasis, cell migration, and cytoskeletal remodeling [55], and the alteration of membrane skeleton stability may further lead to abnormal sodium current, interfering with the contractility phenotype and tension regulation of smooth muscle cells, and subsequently affecting the functional stability of smooth muscle cells [56]. Meanwhile, abnormal sodium channel function has been confirmed to alter cell migration ability [53], the alteration of migration ability may affect the repair and remodeling process of smooth muscle cells at the injury site, indirectly influencing the balance of extracellular matrix deposition and degradation[57, 58]. Animal experiments also show that sodium channel dysfunction can accelerate aortic degeneration [59],which provides indirect evidence for the clinical findings of this study. It can be seen from this that SCN4B may affect the contractile function of smooth muscle cells by regulating ion homeostasis and participate in the remodeling process of the vascular wall by regulating cell migration, thereby playing a role in maintaining the structural integrity of the aortic media. In addition, exercise training has been proven to improve muscle function by regulating the expression of ion channels and electrophysiological characteristics [60],so it is speculated that moderate exercise may maintain the ionic homeostasis and mechanical sensing ability of vascular smooth muscle by up-regulating the expression of SCN4B or enhancing its stability, thereby protecting the integrity of the aortic wall. However, its specific regulatory mechanism in the context of exercise intervention still needs to be further studied and verified through vascular smooth muscle cell-specific models.

It is worth noting that both SCN4B and ABCA3 are involved in eukaryotic ribosome synthesis pathway. Recent studies have shown that ribosome synthesis abnormalities are closely related to the occurrence and development of various cardiovascular diseases[61]. In TAAD, these two key genes may influence the synthesis efficiency of key proteins by co-regulating ribosome function, thus interfering with the proliferation, differentiation and repair ability of VSMC, ultimately compromising aortic wall integrity.

The results of increased M0 macrophage, neutrophil and monocyte infiltration and decreased M1 macrophage and naive B cells in TAAD group in this study are consistent with a study that used both immunohistochemistry and single cell sequencing techniques to analyze in detail the composition of immune cells in the aortic wall of TAAD patients [62]. It was found that the aggregation of pro-inflammatory cells such as M0 macrophages and neutrophils can release a large number of inflammatory mediators, such as cells. Factors and proteases lead to increased inflammation in the blood vessel wall, degradation of elastic fibers, and promotion of dissection formation. Conversely, reduced M1 macrophages and naive B cells may compromise the immune defense and tissue repair ability, impairing the ability to counteract inflammatory damage or maintain vascular homeostasis[63].

Notably, this study found that the expression levels of ABCA3 and SCN4B were significantly associated with the infiltration degree of specific immune cell subsets, suggesting that these two motion-related genes may be involved in the pathological process of TAAD by regulating immune responses. Specifically, the expression of ABCA3 was significantly negatively correlated with neutrophil infiltration. Studies have shown that neutrophils play a significant role in the inflammatory response of the aortic wall by releasing neutrophil extracellular traps and various proteases [64, 65]. Based on this, we speculate that ABCA3 may exert a protective effect on the progression of TAAD by regulating the activity of neutrophil receptors and inhibiting their excessive response to pro-inflammatory signals. On the other hand, The expression of SCN4B shows a positive correlation with the quantity of immature B cells, a result that aligns with the established role of ion channels in regulating immune cell function[66]. Studies have shown that the sodium channel subunit may affect the proliferation and differentiation process of B cells by regulating the B-cell receptor signaling pathway [67]. In view of this, in the pathological environment of TAAD, we speculate that the down-regulation of SCN4B expression may lead to B cell dysfunction through sodium channel regulation, and B cell exhaustion has been confirmed to promote the infiltration of immunosuppressive cells in the aortic wall [68]. Based on these pieces of evidence, we speculate that SCN4B may play a role in inhibiting pathological immune cell infiltration and protecting the structural integrity of the aortic wall by maintaining B-cell homeostasis. These findings collectively reveal a new mechanism by which exercise-related genes may involve immune responses in the pathological process of TAAD. This discovery provides a new immunological perspective for understanding the protective effect of exercise in TAAD, but its specific mechanism of action still needs to be further verified through subsequent experiments. The lncRNA-miRNA-mRNA regulatory axis identified in this study represents a novel therapeutic target for TAAD intervention. Our findings demonstrate that the non-coding RNA regulatory networks centered on ABCA3 and SCN4B play crucial roles in TAAD pathogenesis. This observation is supported by a study [69] which established that lncRNAs and miRNAs can precisely regulate gene expression through sponge mechanisms or by directly binding mRNA by constructing non-coding RNA regulatory networks associated with cardiovascular diseases. The regulation of ABCA3 by hsa-miR-1343-3p and XIST, as well as the regulation of SCN4B by hsa-miR- 4770 and MALAT1, may serve as promising therapeutic entry points for TAAD management. By regulating the expression of these non-coding RNAs, it is hoped that normal levels of ABCA3 and SCN4B expression will be restored, thereby improving disease progression. For example, designing antagonists or mimetics that target specific miRNAs, or regulating lncRNA expression through gene editing techniques, may be new strategies for future TAAD treatments.

Molecular docking analysis in this study suggests that zonisamide and MRS1097 may respectively have good binding properties with SCN4B and ABCA3. This discovery provides new ideas for exploring drug treatment strategies for TAAD. From the perspective of drug mechanism of action, zonisamide, as an antiepileptic drug used in clinical practice [70], has pharmacological effects including the regulation of voltage-gated sodium channels [71]. It is worth noting that existing studies have shown that sodium channel modulators have certain potential in the treatment of cardiovascular diseases, especially in regulating vascular tension, which may play an important role [72]. This suggests that zonisamide may play a regulatory role in the pathological process of TAAD by regulating the function of sodium channels. On the other hand, as an A3 adenosine receptor selective antagonist [73], the potential interaction between MRS1097 and ABCA3 is worthy of in-depth exploration. The adenosine signaling pathway plays a significant role in inflammatory regulation [74], and the inflammatory response of the aortic wall is an important driving factor for the occurrence and development of TAAD [46]. We speculate that MRS1097 may exert a protective effect in TAAD by regulating inflammation. These findings offer new possible directions for the drug intervention strategies of TAAD. However, it should be pointed out that the specific effects and mechanisms of action of these drugs in the treatment of TAAD still need to be verified through further experimental research. Future research should focus on examining the effects of these drugs on the structure and function of the aortic wall in vascular smooth muscle cell models and animal models. At the same time, it is necessary to clarify their direct interaction with target molecules, so as to provide more sufficient experimental evidence for the clinical treatment of TAAD.

In conclusion, based on the analysis results of this study, ABCA3 and SCN4B have demonstrated potential value as diagnostic biomarkers and therapeutic targets for TAAD. This study has several limitations: Firstly, in terms of biomarker screening strategies, based on externally predefined “sports-related genomes” may introduce selection bias, and the functional definition of this genome itself is relatively broad. Secondly, although this study identified exercise-related genes associated with TAAD, it did not directly prove that exercise prevents TAAD by regulating the expression of these genes. The relationship between the two remains at the correlation level at present, and there is a lack of experimental evidence that exercise intervention directly regulates the expression of ABCA3 and SCN4B in vascular tissues. Meanwhile, the main findings, including the correlation of circadian rhythm pathways, all originated from bioinformatics correlation analysis and lacked experimental causal evidence. Specifically, the specific mechanisms of action of ABCA3 and SCN4B in TAAD, their regulatory relationships with circadian rhythms and other pathways (such as the impact of clock gene perturbation on their expression), as well as the drug binding efficacy predicted by molecular docking, all need to be verified through cell and animal models, in vitro binding experiments, and in vivo evaluations. Furthermore, the current analysis fails to integrate multi-omics data to systematically analyze the gene interaction network, and the clinical sample size also needs to be expanded to confirm the universality of the markers. In response to the above issues, future research will expand the sample size, adopt unbiased whole transcriptome screening to verify the current findings, combine functional experiments to establish the causal role of core genes in in vivo and in vitro models, and directly explore the impact of exercise on the expression of these genes and its potential role in the prevention of TAAD through exercise intervention models. At the same time, multi-omics and single-cell technologies are utilized to deeply clarify its regulatory network, and ultimately its clinical transformation and application are promoted through large-scale prospective cohort studies.

## 5. Conclusions

Through multidimensional bioinformatics analysis, the exercise-associated key genes ABCA3 and SCN4B were identified for the first time in TAAD, revealing their multifuctiongal roles in circadian rhythm regulation, immune homeostasis, and cardiovascular function modulation, and predicting potential intervention drugs. The results not only advance our understanding of TAAD’s early diagnostic biomarkers and exercise-modulated pathological mechanisms, but also pave the way for developing precision medicine strategies targeting molecular networks.

## Acknowledgments

We would like to express our sincere gratitude to all individuals and organizations who supported and assisted us throughout this research. Special thanks to Yue Zhang for her technical support. In conclusion, we extend our thanks to everyone who has supported and assisted us along the way. Without your support, this research would not have been possible.

## Supporting information

S1 Table. Metadata of the three motion-related HP gene sets retrieved from MSigDB, along with the merged and deduplicated list of 176 unique genes (Gene_Sets_Summary).

